# Puf4 Mediates Post-transcriptional Regulation of Caspofungin Resistance in *Cryptococcus neoformans*

**DOI:** 10.1101/2020.02.10.943191

**Authors:** Murat C. Kalem, Harini Subbiah, Jay Leipheimer, Virginia E. Glazier, John C. Panepinto

## Abstract

Echinocandins have been on the market for 20 years, yet they are the newest class of antifungal drugs. The human fungal pathogen *Cryptococcus neoformans* is intrinsically resistant to the echinocandin antifungal drug caspofungin, which targets the *β*-1,3-glucan synthase encoded by the *FKS1*. Analysis of a *C. neoformans puf4*Δ mutant, lacking the pumilio/FBF RNA binding protein family member Puf4, revealed exacerbated caspofungin resistance. In contrast, overexpression of *PUF4* resulted in caspofungin sensitivity. The *FKS1* mRNA contains three Puf4-binding elements (PBEs) in its 5’ untranslated region. Puf4 binds with specificity to this region of the *FKS1*. The *FKS1* mRNA was destabilized in the *puf4*Δ mutant, and the abundance of the *FKS1* mRNA was reduced compared to wild type, suggesting that Puf4 is a positive regulator *FKS1* mRNA stability. In addition to *FKS1*, the abundance of additional cell wall biosynthesis genes, including chitin synthases (*CHS3*, *CHS4*, *CHS6*) and deacetylases (*CDA1*, *CDA2*, *CDA3*) as well as a *β*-1,6-glucan synthase gene (*SKN1*) was regulated by Puf4 during a caspofungin time course. The use of fluorescent dyes to quantify cell wall components revealed that the *puf4*Δ mutant had increased chitin content, suggesting a cell wall composition that is less reliant on *β*-1,3-glucan. Overall, our findings suggest a mechanism by which caspofungin resistance, and more broadly, cell wall biogenesis, is regulated post-transcriptionally by Puf4.

**Importance:** *Cryptococcus neoformans* is an environmental fungus that causes pulmonary and central nervous system infections. It is also responsible for 15% of AIDS-related deaths. A major contributor to the high morbidity and mortality statistics is the lack of safe and effective antifungal therapies, especially in resource-poor settings. Yet, antifungal drug development has stalled in the pharmaceutical industry. Therefore, it is of importance to understand the mechanism by which *C. neoformans* is resistant to caspofungin in order to design adjunctive therapies to potentiate its activity toward this important pathogen.

## Introduction

Invasive deep mycoses primarily impact immunocompromised populations causing high rates of morbidity and mortality (1, 2). The pathogenic fungus *Cryptococcus neoformans* is the causative agent of fatal meningitis most often in patients with defects in cell mediated immunity, including transplant recipients and those living with HIV/AIDS (3–6). *C. neoformans* is responsible of the 15% of AIDS-related deaths (6). Treatment of cryptococcosis is difficult and therapeutic options are limited. Even the best combination treatment using Amphotericin B with 5-fluorocytosine (5-FC) is not well tolerated, and 5-FC is often unavailable in resource-poor areas (7). Some of the largest clinal challenges to invasive fungal infections are poor efficacy of the drugs, emerging resistance issues, narrow selection and availability of antifungals especially in the areas where needed the most (8, 9).

Another antifungal agent, fluconazole, is largely ineffective as the first-line therapy since it lacks effective fungicidal activity against *C. neoformans in vivo* even at high concentrations and presents resistance issues (10, 11). Echinocandins (such as caspofungin, micafungin and anidulafungin) are the latest class of antifungal drugs approved by the Food and Drug Administration (FDA) that target cell wall biosynthesis. The echinocandins are ineffective against *C. neoformans* due to a high level of intrinsic resistance (12). Echinocandins specifically target the *β*-1,3-glucan synthase encoded by *FKS1,* and *C. neoformans* Fks1 is sensitive to inhibition by the echinocandins, suggesting that the mechanism of intrinsic resistance is not related to biochemical differences in the target itself (13). In other pathogenic fungi, such as *Aspergillus* and *Candida* species, resistance to caspofungin is manifested due to mutations in *FKS1*, but this is not observed in *Cryptococcus* (14, 15). Another mechanism that is discussed regarding caspofungin resistance in pathogenic fungi involves the cell wall remodeling and integrity pathways (14–16). It has been shown that increased cell wall chitin content contributes to caspofungin resistance (17, 18). Additionally, a defect in intracellular drug concentration maintenance due to drug influx and efflux imbalance has been proposed to be a potential mechanism to explain the intrinsic resistance phenomenon (19, 20). Discovering and targeting the regulatory components behind the pathways involved in the intrinsic resistance may result in a combination therapy that potentiates the antifungal activity of caspofungin toward *C. neoformans*.

Calcineurin signaling plays a distinct role in intrinsic caspofungin resistance (21, 22). Caspofungin synergizes with the calcineurin inhibitors FK506 and Cyclosporin A (23). Both the A and B subunits of Calcineurin regulate tolerance to caspofungin (19). Crz1, the transcription factor that is activated through dephosphorylation by calcineurin, translocates to the nucleus following treatment with caspofungin, yet the *crz1*Δ mutant exhibits wild type sensitivity to the drug, suggesting caspofungin resistance is Crz1-independent. (19). Calcineurin functions at the intersection of multiple signaling pathways and interacts with a diversity of proteins involved in calcium signaling, RNA processing, protein synthesis and vesicular trafficking among others (24, 25). Since caspofungin resistance is calcineurin-dependent, yet Crz1-independent, RNA processing targets of calcineurin that may be involved in resistance to caspofungin through post-transcriptional modulation of gene expression are especially of interest. Tight control of not only transcriptional but also post-transcriptional gene regulatory networks in drug resistance phenotypes is underappreciated in fungal pathogens, yet may represent targets for adjunctive therapies to improve the efficacy of drugs.

One of the targets of calcineurin involved in RNA processing is a pumilio-domain and FBF (PUF) domain-containing RNA-binding protein - Puf4 (24). *C. neoformans* Puf4 is homologous to the *Saccharomyces cerevisiae* paralogs Puf4 and Mpt5 (26). Our previous work demonstrated that *C. neoformans* Puf4 recognizes the Mpt5 binding element in its mRNA targets (27). It has been hypothesized that Puf4 may play a role in the regulation of cell wall biosynthesis since the *puf4*Δ mutant is resistant to lysing enzymes, is temperature sensitive at both 37°C and 39°C, and is sensitive to Congo red (28). Our group has previously shown that Puf4 regulates endoplasmic reticulum (ER)-stress through controlling the splicing of a major ER-stress related transcript, *HXL1,* and plays a role in the unfolded protein response pathway of *C. neoformans.* Puf4 also contributes to virulence, as the *puf4*Δ mutant has attenuated virulence compared to wild type in an intravenous murine competition model of cryptococcosis (27).

In this study we demonstrated that Puf4 contributes to the intrinsic caspofungin resistance of *C. neoformans* through post-transcriptional regulation of the mRNA encoding the drug target, Fks1. Puf4 also regulated a number of the cell wall biosynthesis-related mRNAs. This regulation is primarily at the level of mRNA stability and has functional consequences in maintaining cell wall composition.

## Results

### The *puf4*Δ mutant is resistant to caspofungin

Our previous work has implicated the pumilio family RNA binding protein Puf4 in the regulation of ER stress in *C. neoformans* (27). Puf4 is an effector of calcineurin signaling, a pathway known to regulate thermotolerance and cell integrity (24, 28). Given the connection of Puf4 to cell integrity signaling, we assessed the sensitivity of *puf4Δ* mutant to the cell wall perturbing drug caspofungin. We measured caspofungin sensitivity by spot plate analyses, and found that the *puf4*Δ mutant is resistant to caspofungin above the published minimum inhibitory concentration of 16 µg/ml (Fig. 1A). This result suggests that Puf4 is a negative regulator of caspofungin resistance in *C. neoformans,* since its absence presents a hyper-resistant phenotype. When Puf4 was overexpressed with a FLAG-tag (3-copies determined by Southern blot analysis), the hyper-resistance was not only suppressed, but the strain was more sensitive to caspofungin. Single-copy FLAG-Puf4 complementation of the *puf4*Δ mutant restored a wild type resistance phenotype (Fig. 1A). In addition to the spot plate analyses, growth analysis using liquid cultures in a kinetic plate reader assay in the presence of 8 µg/ml caspofungin showed similar trends to the phenotypes observed in spot plates (Fig. 1B). Growth was not permissible in the presence of 16 µg/ml caspofungin in liquid culture for both wild type and the mutant cells suggesting that the action of caspofungin may be influenced by the environmental constraints in different culture conditions (data not shown).

**Figure 1.**
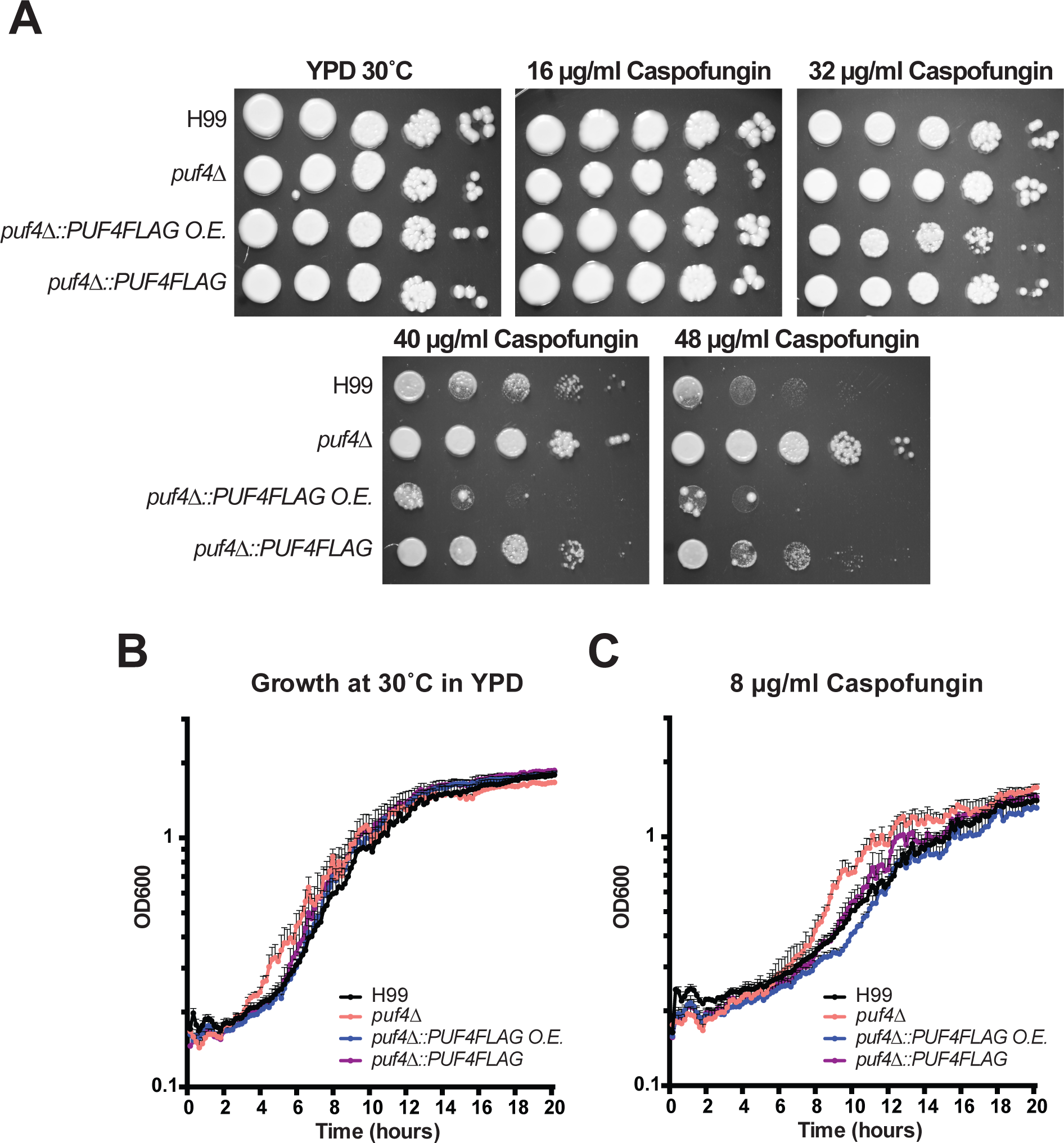
The *puf4*Δ mutant is resistant to caspofungin. (A) Spot plate analysis. The indicated strains were diluted to an OD_600_ of 1.0 and five, 1:10 serial dilutions were spotted on agar plates containing 0, 16, 32, 40, and 48 µg/ml caspofungin. Plates were incubated at 30°C for 3 days and photographed. (B-C) Growth assay. The indicated strains were diluted to an OD_600_ of 0.3 and mixed 1:1 with either fresh YPD or YPD containing caspofungin in a 96 well-plate. Plates were incubated and at 30°C for 20 hours while shaking. OD_600_ was measured every 10 minutes.

### Puf4 directly binds and stabilizes the *FKS1* mRNA

The target of caspofungin, *β*-1,3-glucan synthase, is encoded by the *FKS1* gene. We searched the *FKS1* mRNA sequence for a potential Puf4 binding element. Our previous work suggests that the Puf4 binding element in *C. neoformans* is homologous to that of *S. cerevisiae* Mpt5, including the invariant initiating UGUGA followed by a four-nucleotide spacer sequence and terminating UA. We found that the *FKS1* mRNA contains two Puf4 binding elements in its 5’ UTR (Fig 2A). To determine if Puf4 can directly interact with its consensus element(s) in the 5’UTR of the *FKS1* mRNA we performed electrophoretic mobility shift assay (EMSA). We synthesized a 50-base long fluorescently labeled (TYE705 infrared label) RNA oligonucleotide that span the *FKS1* 5’ UTR containing the Puf4-binding elements (Table 2). Incubation of the fluorescent oligonucleotide with the wild type cell lysate resulted a shift that was competed with the unlabeled oligonucleotide. A mutant competitor, in which the **UGUA**NNNN**UA** motif was replaced by adenines, was unable to compete for binding to the wild type fluorescent oligonucleotide. This suggests that Puf4 binds to the *FKS1* mRNA through sequence-specific recognition of the Puf4 binding elements in the 5’ UTR (Fig. 2B). Incubation of the fluorescent oligonucleotide with the *puf4*Δ mutant lysate has the absence a shift that was observed with the Puf4-FLAG cell lysate (Fig 2B, arrow). Because, we detected an interaction between Puf4 and the 5’UTR sequence of *FKS1*, we went on to investigate if loss of Puf4 would alter the abundance or stability of *FKS1* mRNA. *FKS1* mRNA abundance in mid-log grown cultures of *puf4*Δ cells were decreased 20% compared to wild type grown in parallel (Fig. 2C). PUF proteins are known mRNA stability regulators, and so we asked if the reduction in *FKS1* mRNA in the *puf4Δ* mutant was due to destabilization. We performed an mRNA stability time-course following transcription shut-off and found that the *FKS1* mRNA was destabilized in the absence of Puf4 compared to the wild type (Fig 2D).

**Figure 2.**
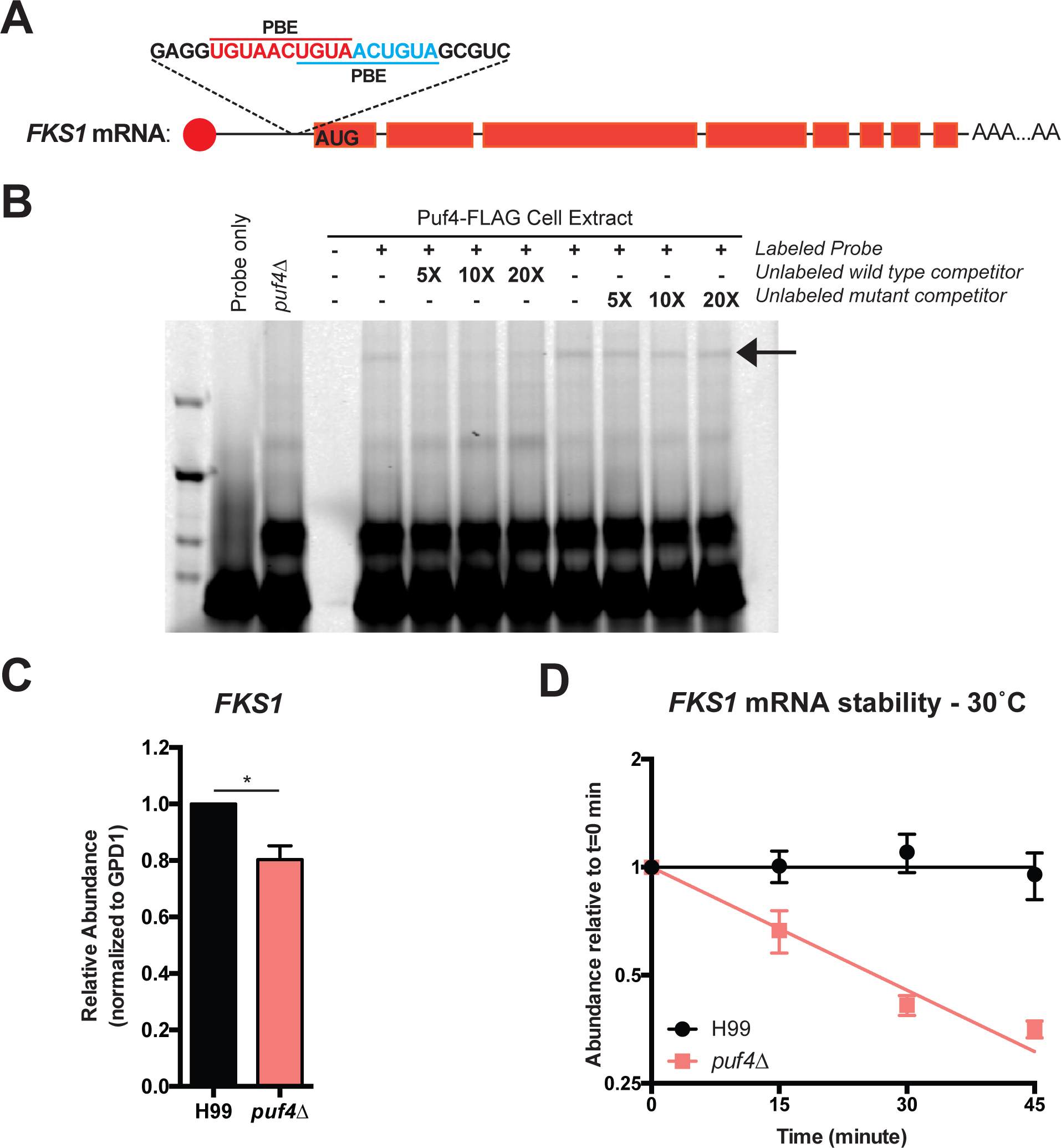
Puf4 directly binds and stabilizes the *FKS1* mRNA. (A) The *FKS1* mRNA contains Puf4 binding elements (PBE) in its 5’ UTR - UGUANNNNUA. (B) Puf4 binds to *FKS1*. Electrophoretic mobility shift assay was performed using a fluorescently labeled synthetic RNA oligonucleotide designed for the *FKS1* 5’UTR that contain the PBEs. Unlabeled mutant (contains mutated PBEs) and unlabeled wild type probes were used as controls for sequence specificity. *puf4*Δ was included as a control to show binding is absent when Puf4 is not present. A representative gel image is shown, n=3. (C) The *FKS1* is downregulated in the *puf4*Δ. The *FKS1* mRNA abundance in mid-log samples grown at 30°C was determined using RT-qPCR with *GPD1* as the normalization gene. 3 replicates were plotted and unpaired t-test with Welch’s correction was performed, *: p<0.05. (D) The *FKS1* is destabilized in the *puf4*Δ mutant. *FKS1* abundance was determined using RT-qPCR following transcription shut-off to determine the decay kinetics. *GPD1* was utilized as the control for normalization. 3 replicates were plotted and differences among two strains were analyzed using one-phase exponential decay analysis. Error bars show SEM.

**Table 2.**
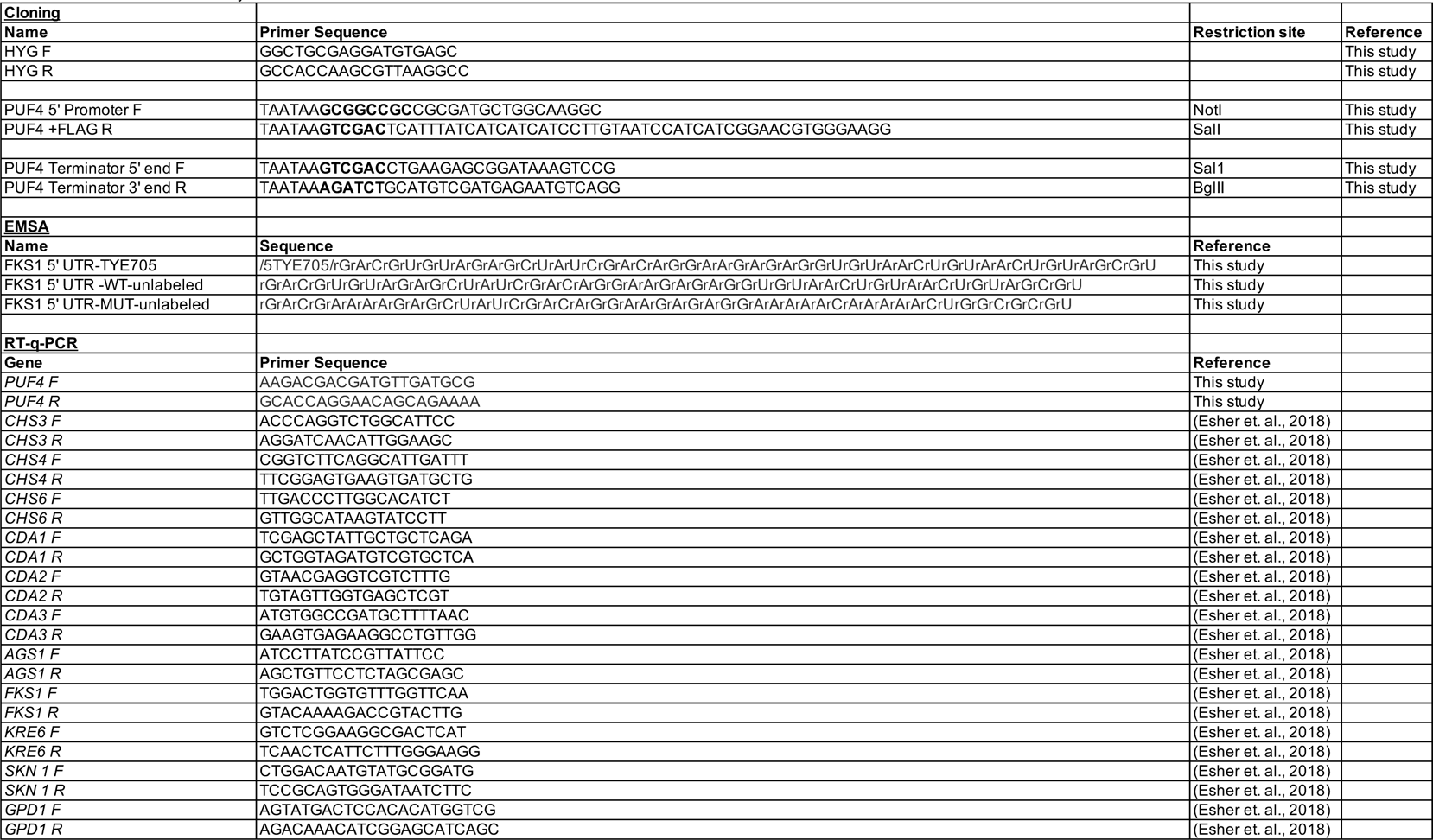
Primers used in this study

### Puf4 protein expression is decreased following caspofungin treatment

Because Puf4 is a regulator of the *FKS1* mRNA, we asked if inhibition of Fks1 by caspofungin treatment would alter the abundance of *PUF4* mRNA or Puf4 protein. First, we performed a caspofungin time course in which we treated wild type cultures grown to mid-log phase with 16µg/ml caspofungin for 60 minutes and collected cells every 15 minutes. We quantified the *PUF4* transcript levels, and found out that the *PUF4* transcript levels are unchanged throughout time-course (Fig 3A). Then we utilized the Puf4-FLAG strain to investigate if Puf4 protein levels are changed following treatment with caspofungin using immunoblotting. We found that Puf4 protein levels are drastically decreased following treatment with caspofungin (Fig 3B and 3C). Since the absence of Puf4 causes a hyper-resistant phenotype (as shown in Fig. 1A), we speculate that the downregulation of Puf4 protein levels may be a contributing event to the intrinsic resistance.

**Figure 3.**
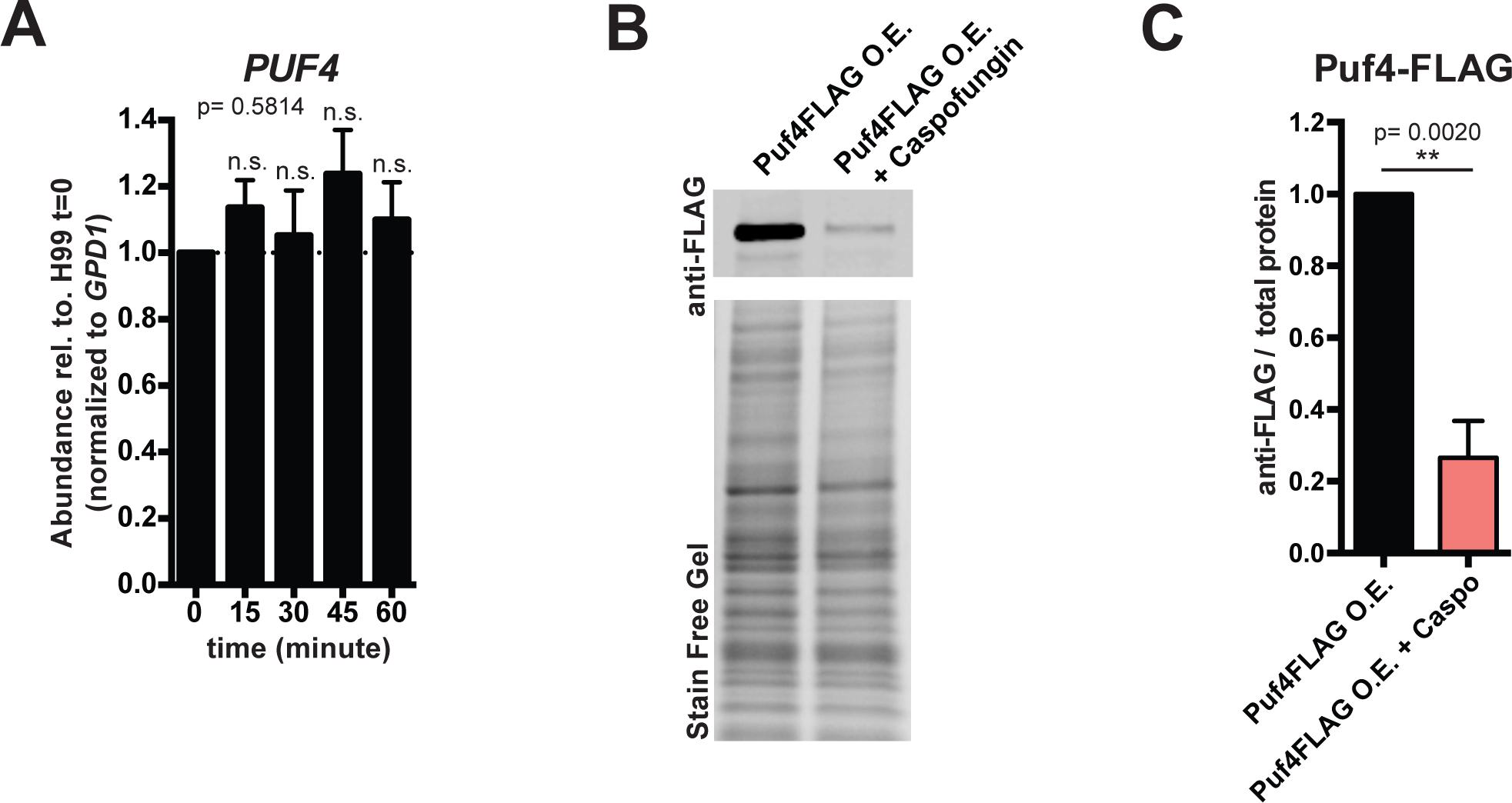
Puf4 protein expression is decreased following caspofungin treatment. (A) *PUF4* transcript levels are unchanged during a 60-minute caspofungin time course. Cells were grown at 30°C and treated with caspofungin. The abundance of *FKS1* mRNA was determined in samples collected every 15 minutes using RT-qPCR with *GPD1* as the normalization gene. 3 replicates were plotted and one-way-ANOVA using Tukey’s test was performed, p=0.5814 (B) Puf4 protein levels are decreased following 60-minute caspofungin treatment. Puf4-FLAG strain (*puf4Δ* expressing *PUF4-FLAG* C-terminal fusion) was grown to mid-log and treated with caspofungin for 1 hour. SDS-PAGE followed by immunoblotting using anti-flag antibody is shown. (C) Anti-flag signal is normalized to total protein signal from the stain-free gel. 5 replicates were plotted and unpaired t-test with Welch’s correction was performed, **: p<0.01. Error bars show SEM.

### Puf4-dependent *FKS1* regulation correlates with reduced β-1,3-glucan staining in the *puf4*Δ mutant

We next investigated the *FKS1* mRNA levels following treatment with caspofungin to determine if inhibition of the enzyme activates a feedback mechanism to upregulate expression of *FKS1*., The *FKS1* mRNA abundance is upregulated in wild type cells at 45 minutes post-caspofungin treatment compared to 30 minutes (Fig 4A). Conversely, the *puf4*Δ mutant cells have decreased levels of *FKS1* at 30 minutes post-caspofungin treatment, which is restored to the basal levels at later time points (Fig. 4A). These different trends in transcript abundance following caspofungin treatment may suggest that the cell wall *β*-1,3-glucan levels may differ between wild type and the *puf4*Δ cells.

**Figure 4.**
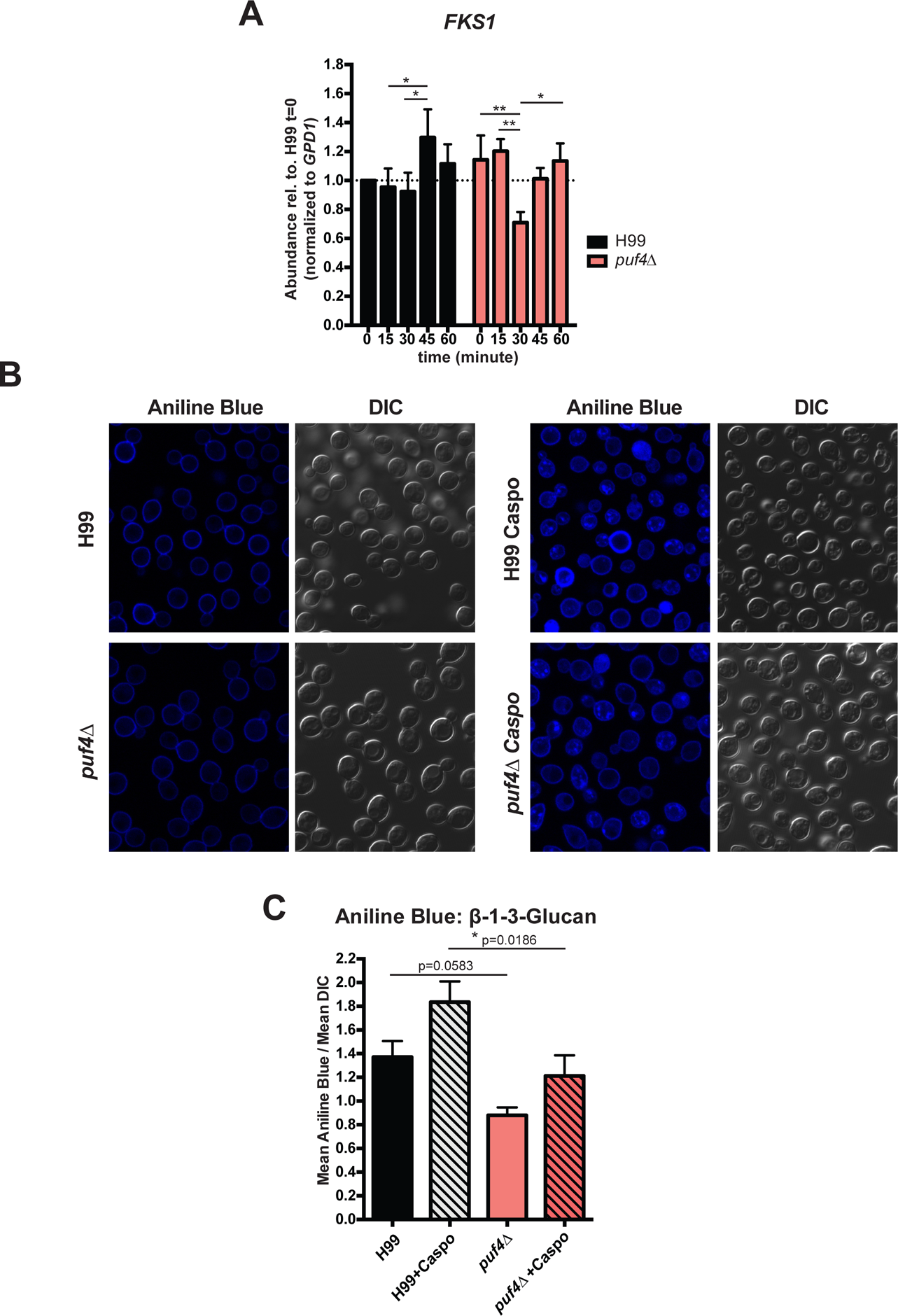
Puf4-dependent *FKS1* regulation correlates with reduced β-1,3-glucan staining. (A) *FKS1* abundance is regulated by Puf4 during a 60-minute caspofungin time course. Cells were grown at 30°C and treated with caspofungin. The abundance of *FKS1* mRNA was determined in samples collected every 15 minutes using RT-qPCR with *GPD1* as the normalization gene. 3 replicates were plotted and one-way-ANOVA using Tukey’s test was performed. (B) The *puf4*Δ mutant has decreased levels of *β*-1,3-glucan. Cells were grown to mid-log at 30°C and stained with Aniline Blue to detect *β*-1,3-glucan. Representative Aniline Blue staining images for each strain is shown to assess both levels and localization. (C) Microscopy images were quantified on Fiji and Aniline Blue signal was normalized to DIC signal. At least 5 fields from 3 biological replicates were plotted and one-way-ANOVA followed by Dunn’s multiple comparisons test was performed. *: p<0.05, **: p<0.01 Error bars show SEM.

To investigate the functional consequences of the trends we observed at the transcript level, we utilized aniline blue staining to investigate the *β*-1,3-glucan levels in the cell wall of the wild type and the *puf4*Δ cells. Aniline blue specifically binds to *β*-1,3-glucan (29). Following growth to mid-log phase, both strains were treated with caspofungin for 60 minutes and stained with aniline blue. Fluorescence microscopy revealed that aniline blue staining mainly localized to the cell wall in the mid-logarithmic stage cells. In addition to the cell wall staining pattern, staining pattern of the caspofungin treated cells also contained an intracellular punctate pattern (Fig. 4B). Quantification of the microscopy images showed that the *puf4*Δ cells had 30% less *β*-1,3-glucan compared to the wild type cells both at the mid-logarithmic stage and following treatment with caspofungin (Fig. 4C). We concluded that the post-transcriptional regulation of the *FKS1* mRNA by Puf4 directly affects cellular *β*-1,3-glucan levels.

### Deletion of *PUF4* leads to dysregulation of cell wall biosynthesis genes

A recent study revealed that multiple cell wall genes are influenced by caspofungin treatment (19). We assessed the same panel of mRNAs for the presence of a putative Puf4 binding element, and found that several caspofungin-sensitive genes contain Puf4 binding elements (Table 1). These include genes that encode chitin synthases, chitin deacetylases, *α*-glucan and *β*-glucan synthases.

**Table 1.** List of Puf4 binding element containing cell wall biosynthesis related genes. **UGUA**NNNN**UA** motif was searched in target genes using FIMO (MEME-suite version 5.1.1.). Results were manually confirmed and location of the motifs were identified. **\*** Indicates the genes selected for further mRNA stability analysis.

| Gene | Gene ID | Start | End | p-value | Location |
| --- | --- | --- | --- | --- | --- |
| ***CHS3\**** | CNAG_05581 | 242 | 251 | 0.000262 | 5' UTR |
| ***CHS4\**** | CNAG_00546 | 3257 | 3266 | 0.00013 | Exon |
|  |  | 3516 | 3525 | 0.000203 | Exon |
| ***CHS7*** | CNAG_02217 | 88 | 97 | 6.59E-05 | 5' UTR |
|  |  | 3264 | 3273 | 6.59E-05 | Exon |
| ***CHS8*** | CNAG_07499 | 1479 | 1488 | 7.96E-05 | Exon |
| ***CDA3\**** | CNAG_01239 | 1916 | 1925 | 7.96E-05 | 3' UTR |
| ***FKS1\**** | CNAG_06508 | 281 | 290 | 7.96E-05 | 5' UTR |
|  |  | 307 | 316 | 0.000162 | 5' UTR |
|  |  | 313 | 322 | 0.000162 | 5' UTR |
| ***SKN1*** | CNAG_00897 | 2471 | 2480 | 0.000122 | 3' UTR |
| ***AGS1\**** | CNAG_03120 | 202 | 211 | 0.000107 | 5' UTR |

We next asked if the caspofungin-responsiveness of these genes was dependent on Puf4. Following growth to the mid-log stage, we challenged both wild type and the *puf4*Δ cells with 16 µg/ml caspofungin over a 60-minute time course and investigated the changes in the transcript abundance of genes involved in cell wall biosynthesis. We found that *CHS3* (Chitin Synthase 3) is downregulated following caspofungin treatment in the *puf4*Δ cells compared to wild type (Fig 5A). Conversely, *CHS4* and *CHS6* are found to be upregulated in the *puf4*Δ mutant compared to wild type (Fig 5B and 5C). Elevated cell wall chitin content is shown to reduce susceptibility to caspofungin in Candida species (18). Therefore, upregulation of chitin synthase genes in the *puf4*Δ mutant may also contribute to the resistance phenotype in Cryptococcus. The synthesis of chitosan from chitin is catalyzed by the chitin deacetylases and chitosan is necessary for the integrity of the cell wall (30). During the caspofungin time course, we found out that *CDA1* (chitin deacetylase 1) is upregulated in the *puf4*Δ mutant. On the contrary, *CDA2* and *CDA3* were found to be downregulated in the *puf4*Δ mutant (Fig 5D-F). The CDA3 is the only chitin deacetylase gene that contain a PBE (Table 1). Of note, Cda1 is the major chitin deacetylase and the only chitin deacetylase that is necessary for virulence (31).

**Figure 5.**
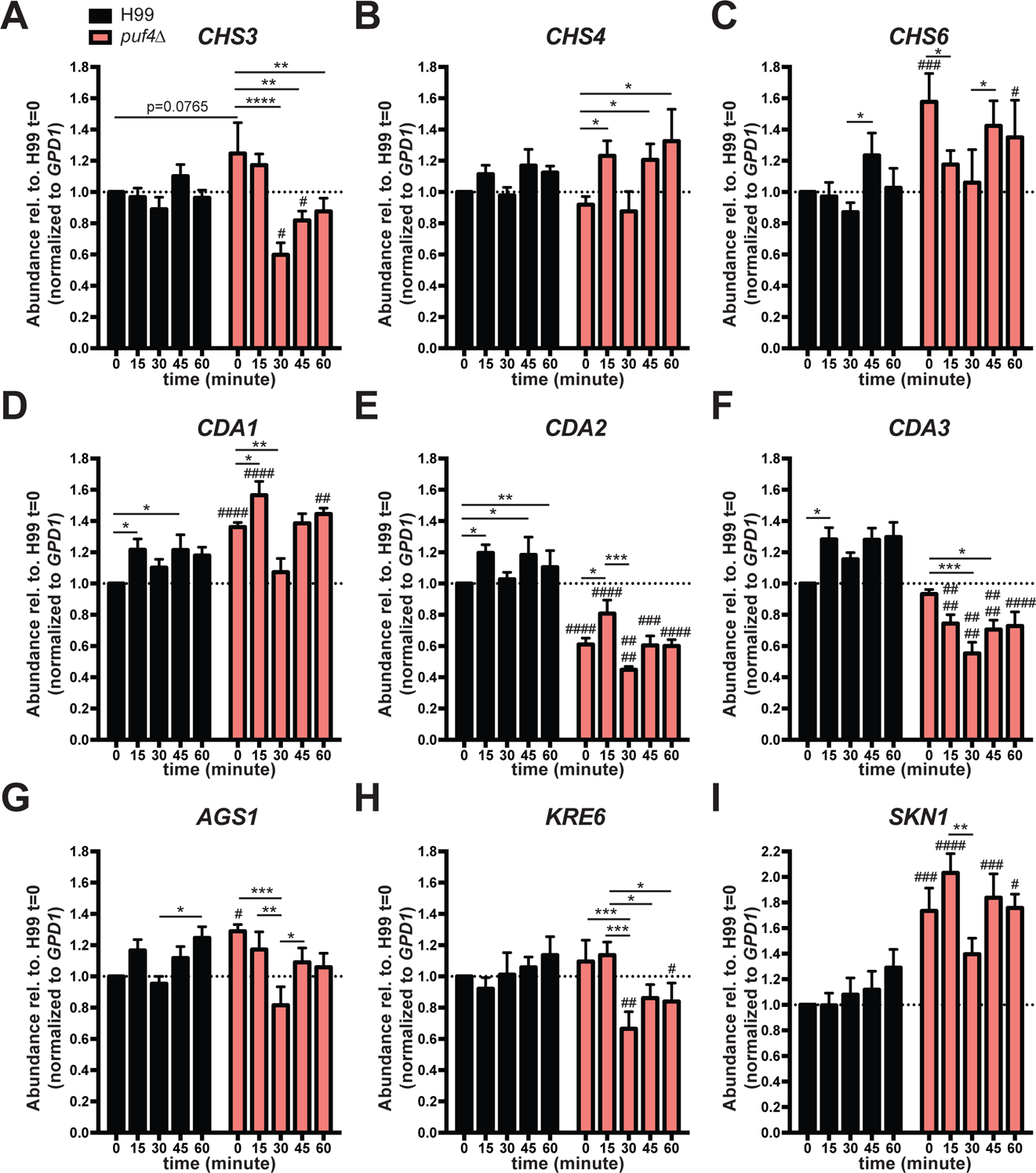
Deletion of *PUF4* leads to dysregulation of certain cell wall biosynthesis genes. The mRNA abundance of select cell wall biosynthesis genes was determined during a 60-minute caspofungin time course by collecting samples every 15 minutes and determining abundance by RT-qPCR using *GPD1* as a normalization gene. (A) *CHS3,* (B) *CHS4,* (C) *CHS6,* (D) *CDA1,* (E) *CDA2,* (F) *CDA3,* (G) *AGS1,* (H) *KRE6,* and (I) *SKN1*. 3 biological replicates with 2 technical replicates were plotted and two-way ANOVA was used to determine statistical significance. ‘#’ denotes comparison between wild type and *puf4*Δ while ‘*’ denotes comparison between indicated time points within a strain. #/*: p<0.05, ##/**: p<0.01, ###/*** p<0.001 and ####/****: p<0.0001. Error bars show SEM.

Lastly, we looked at the regulation of *α*-glucan and *β*-glucan genes during the caspofungin time-course. We have found that *AGS1* (*α*-glucan synthase 1) was present at a slightly higher abundance in the *puf4*Δ compared to the wild type at the basal levels. Which then decreased significantly at the 30 minutes time point. The *AGS1* contains a PBE at its 5’UTR, and may be a Puf4 target (Table 1). The *β*-1,6-glucan synthase genes *KRE6* and *SKN1* showed opposite trends. Both wild type and the *puf4*Δ cells had comparable levels of *KRE6* at t=0 minutes, yet the *puf4*Δ cells had a significantly decreasing trend of *KRE6* abundance during the caspofungin time-course. Another *β*-1,6-glucan synthase gene *SKN1* was upregulated in the *puf4*Δ compared to wild type and remained upregulated throughout the 60 minutes (Fig 5G-I). The PBE element that is present in the *SKN1* 3’ UTR compared to KRE6, which does not have a PBE, may explain the opposite trends in post-transcriptional gene regulation.

Our quantitative analysis of the mRNAs involved in cell wall biosynthesis showed that Puf4 plays a regulatory role in the fate of these mRNAs and modulate their abundances during caspofungin treatment.

### Puf4 stabilizes cell wall biosynthesis genes involved in chitin and α-glucan synthesis

To gain more mechanistic insight on how Puf4 may control the cell wall biosynthesis related transcript abundances, we investigated the mRNA stability of the same transcripts. mRNA stability is a crucial step in transcriptome remodeling to adapt to various environmental and compound stressors (32). Therefore, we hypothesized that Puf4 may modulate cell wall biosynthesis genes post-transcriptionally at the mRNA stability level.

We found that *CHS3* and *CHS4* were destabilized to a great degree in the *puf4*Δ mutant, whereas *CDA3*, *AGS1* and *CDA1* exhibited a slight reduction in stability in the absence of Puf4. Unlike other genes we investigated, *CDA1* does not contain a PBE and was included as a negative control. Even though we expected *CDA1* stability to be like wild type, it too was slightly destabilized (Fig. 6). Our results show that Puf4-mediated post-transcriptional gene regulation in mRNA stability may be crucial for cell wall remodeling that contributes to the caspofungin resistance.

**Figure 6.**
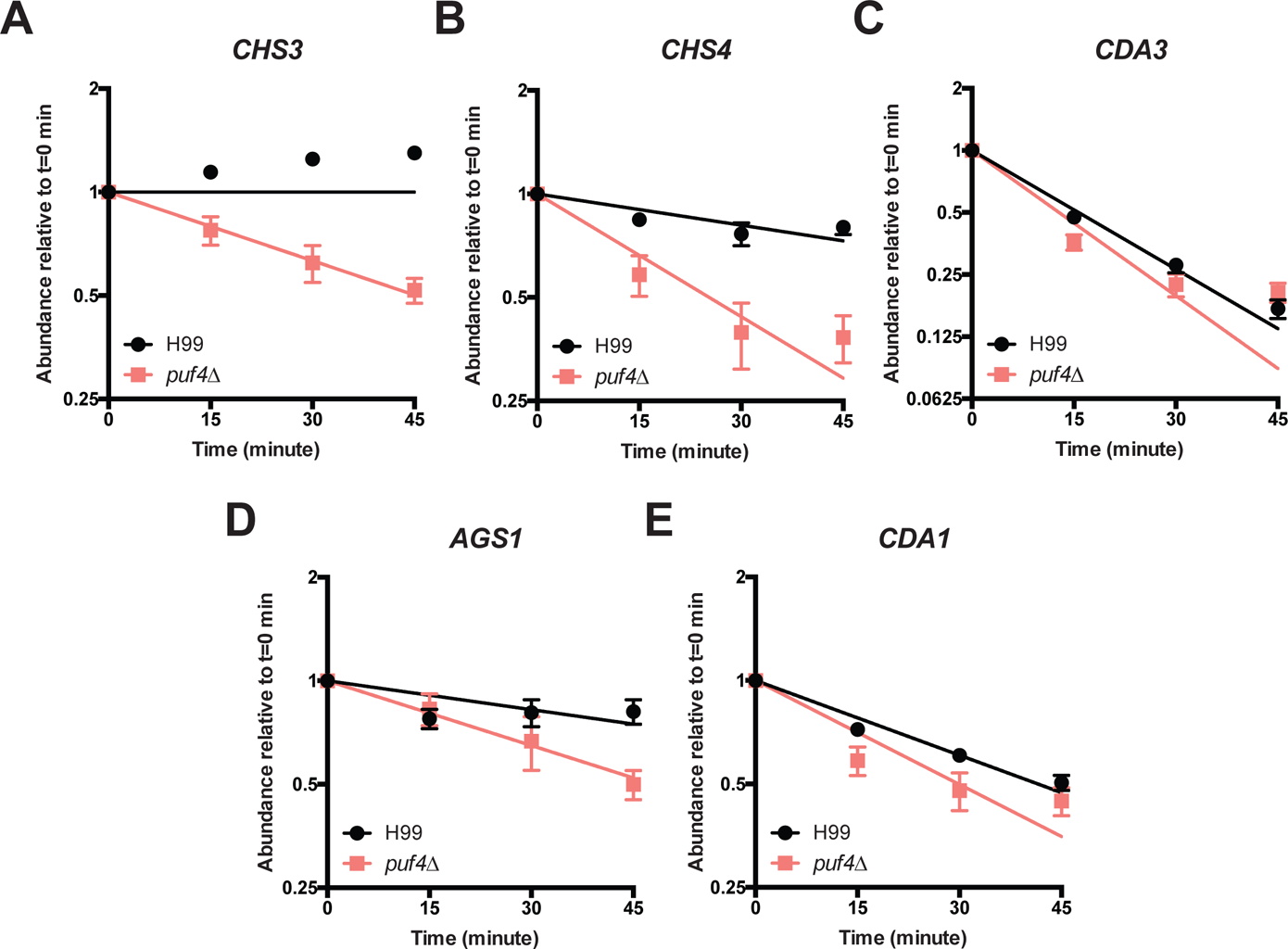
Puf4 stabilizes cell wall biosynthesis genes involved in chitin and α-glucan synthesis. (A) *CHS3*, (B) *CHS4*, (C) *CDA3*, (D) *AGS1*, (E) *CDA1* transcript abundance were determined using RT-qPCR following transcription-shutoff to determine the decay kinetics. *GPD1* was utilized as the control for normalization. 15 minutes post-transcription shut-off was denoted as t=0. 3 replicates were plotted and differences among two strains were analyzed using one-phase exponential decay analysis. Error bars show SEM.

### Caspofungin treatment leads to increased cell wall chitin which is exacerbated in the ***puf4*Δ mutant.**

Since we have shown that Puf4 regulates cell wall biosynthesis related transcript abundances and their mRNA stability, we further investigated the functional consequences of this Puf4 loss by assessing cell wall chitin and chitooligomer content using calcofluor and wheat germ agglutinin staining, respectively. Microscopy (Fig 7A) and flow cytometry analysis (Fig 7B-C) of the chitin content using calcofluor dye showed that the cell wall chitin content increases following caspofungin treatment in both wild type and the *puf4*Δ cells. Importantly, the *puf4*Δ cells had significantly more chitin following caspofungin treatment compared to that of wild type cells. In other pathogenic fungi such as *Candida* species and *Aspergillus* fumigatus, increased chitin content is protective against caspofungin (17, 18).

**Figure 7.**
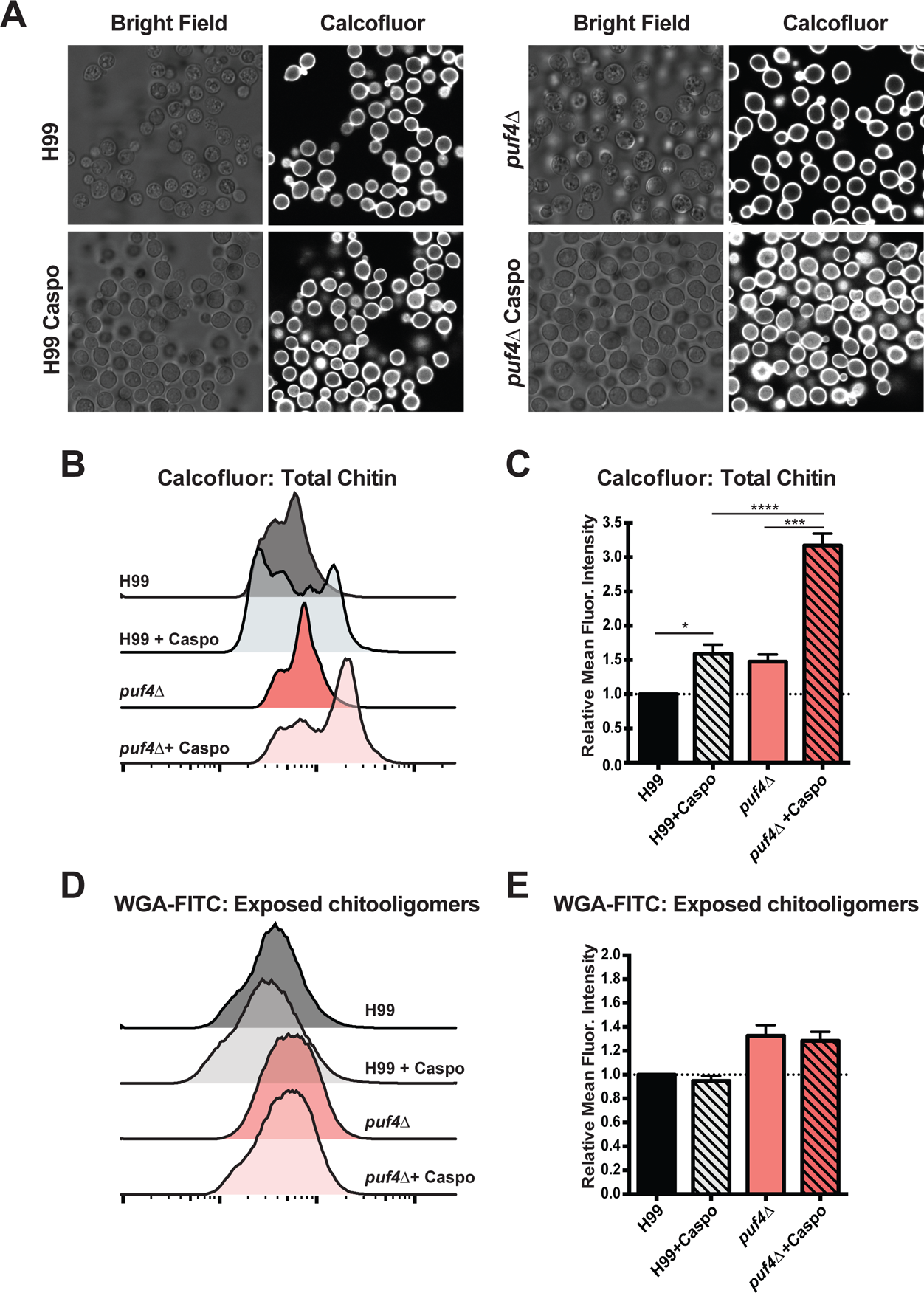
Caspofungin treatment leads to increased cell wall chitin which is exacerbated in the *puf4*Δ mutant. (A-C) Cell wall chitin levels are increased response to caspofungin. Cells were grown to mid-log at 30°C and stained with calcofluor white, then fluorescent intensity (A-B) and staining pattern (C) were determined using flow cytometry and fluorescence microscopy, respectively. (D-E) Levels of exposed chitooligomers did not change significantly. Exposed chitooligomers were stained using wheat germ agglutinin conjugated with FITC. Fluorescent intensity was determined using flow cytometry. For (A) and (D), 3 biological replicates were plotted and one-way-ANOVA followed by Dunn’s multiple comparisons test was performed. *: p<0.05, *** p<0.001 and ****: p<0.0001. Error bars show SEM.

Microbial cultures show molecular and phenotypical heterogeneity that may be important within the scope of antimicrobial resistance (33). Histogram graphs of the calcofluor staining showed that the increase in chitin content is at a sub-population level (Fig 7B). While the majority of the *puf4*Δ cell population shifted to a high chitin phenotype, wild type cells showed a more heterogeneous population. Lastly, we looked at the exposed chitooligomers as another form of chitin-derived structure, and saw a modest increase compared in the *puf4*Δ cells compared to the wild type.

In this study, we have demonstrated that the caspofungin resistance of *C. neoformans* is regulated at the post-transcriptional level through the direct interaction of the mRNA encoding its target, *FKS1*, as well as through the regulation of multiple genes that regulate cell wall composition. Post-transcriptional regulation of cell wall remodeling genes by Puf4 has functional consequences, as the absence of Puf4 results to massive remodeling of the *C. neoformans* cell wall components. Future work will investigate the effect of Puf4 on the translation of these target mRNAs, as well as the mechanism by which Puf4 itself is regulated.

## Discussion

The search for novel antifungal therapies is an ongoing battle in medical mycology, especially with the growing number of fungal outbreaks and emerging drug resistance issues (33, 34). The latest class of antifungals approved by the FDA are the echinocandins, and *Cryptococcus* is intrinsically resistant to this class of antifungals (9, 14, 15, 36). Even though it is crucial to design new therapies, it is also imperative to understand the resistance mechanisms to the existing antifungals to avoid similar scenarios and to design adjunctive therapies to remedy the current resistance issues. In this study, we elucidated the role of post-transcriptional gene regulation in the molecular mechanism of action behind the caspofungin resistance. We discovered that Puf4, a Pumilio domain-containing RNA binding protein, plays a role in the resistance phenotype by stabilizing the mRNAs encoding cell wall biosynthesis genes post-transcriptionally. The functional consequence of this interaction is a change in cell wall composition to a state that is more favorable during caspofungin challenge (Fig. 8). The intrinsic resistance of *C. neoformans* to caspofungin involves multiple signaling pathways (13, 35). For the first time, we have implicated post-transcriptional regulation of cell wall biosynthesis mRNAs, including the mRNA encoding the target of caspofungin, in this intrinsic resistance.

**Figure 8.**
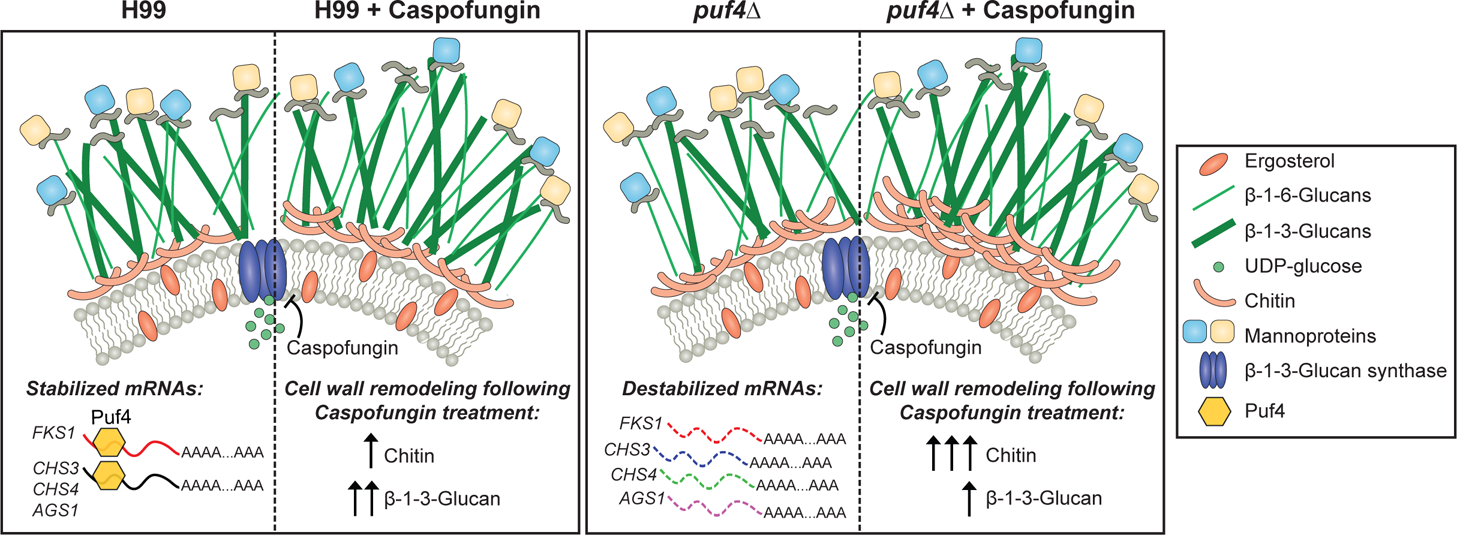
Model: Post-transcriptional regulation of cell wall remodeling by Puf4 is a path to caspofungin resistance in *C. neoformans*. In the wild type cells, *FKS1* is stabilized by Puf4. When wild type cells are treated with caspofungin, we observe increase of cell wall chitin and *β*-1,3-glucan staining intensity. In the *puf4*Δ, *FKS1* as well as other cell wall biosynthesis transcripts are destabilized. Therefore, we observed a different response to caspofungin in the *puf4*Δ involving a robust increase in cell wall chitin content. Graphics modified from (9).

PUF proteins, in addition to their interactions with other signaling proteins, alters mRNA function, and this is often secondary to the translational repression or the inhibition of mRNA decay (37, 38). Binding by Puf4 and other PUF proteins orchestrate mRNA fate-determining processes including stability, splicing, localization and translatability (39, 40). For example, *S. cerevisiae* Puf4p stabilizes the transcripts involved in rRNA processing, and deletion of *PUF4* in *S. cerevisiae* causes defects in translation. Additionally, Puf4p plays a role in the recruitment of mRNAs to the translational machinery (41). In *C. neoformans*, Puf4 appears to play both positive and negative regulatory roles. Puf4 is a positive regulator of the unconventional splicing of the ER-stress transcription factor *HXL1*. In contrast, Puf4 is a negative regulator of the *ALG7* mRNA, which is stabilized in the *puf4Δ* mutant (27). Interestingly, the Puf4 elements in the *HXL1* mRNA, like *FKS1*, are in the 5’ UTR, which may suggest that 5’ UTR Puf4-binding elements exert positive regulatory activity. Stabilization of the 5’ UTR Puf4 binding element containing *CHS3* and *AGS1* in the absence of Puf4 support this claim (Table 1 and Fig. 6). Caspofungin challenge likely requires a reprogramming of gene regulatory networks for adaptation, and Puf4 and other related RNA-binding proteins may be involved in transforming the translating mRNA pool to best respond to the pharmacologically induced stress by caspofungin.

Cell wall maintenance and perturbations in response to drug induced stress have a broad appreciation by the medical mycology (42). For example, the *mar1*Δ mutant exhibits a defect in intracellular trafficking of cell wall synthases and therefore exhibits a cell wall composition that contains elevated exposed chitin and decreased glucan levels. These changes in the cell wall composition and exposure of different carbohydrates play meaningful roles in the immune recognition by the host and activate various signaling events in the host system (43). Another example is the enhanced recognition of the *ccr4*Δ mutant by alveolar macrophages due to increased unmasking of the *β*-1,3-glucan (44). The importance of Ccr4, an mRNA deadenylase, in glucan masking is further evidence that post-transcriptional processes are essential for adaptation to a number of stressors, including caspofungin treatment. In this study, we show that Puf4 is a major regulator of cell wall biogenesis. We report that cell wall biosynthesis genes showed different trends of expression in the *puf4*Δ compared to wild type in caspofungin time-course experiments. We have also shown that *FKS1*, *CHS3* and *CHS4* mRNAs were destabilized in the absence of Puf4. This regulation of certain cell wall biogenesis genes by Puf4 may be a necessary component of cell wall homeostasis under normal growth conditions as well as facilitating rapid changes in cell wall gene expression during adaptation to drug-induced stress. We have predicted that transcripts that have the PBE would have enhanced decay in the *puf4*Δ mutant yet this was not true for all transcripts that carried PBEs. We investigated the stability of *CDA1* as a transcript that did not contain a PBE, yet we observed a modest change in the mRNA stability. In that regard, it must be noted that other RNA-binding proteins may play meaningful regulatory roles in conjunction with Puf4. Additionally, the impact of Puf4 on its targets may be unique and this may be due to the location of the PBE within the transcript, whether it is in the 3’ or 5’ UTR, introns or exons, and may yield to variations in which Puf4 regulates transcript fate. Moving forward, screening additional RNA-binding proteins to understand the post-transcriptional regulatory network controlling cell wall dynamics, as well as investigating the regulatory connections with the cell integrity pathway and calcineurin signaling pathway will elucidate further the response of *C. neoformans* to caspofungin that mitigate its toxicity and promote intrinsic resistance.

The complex cell wall structure of *C. neoformans* protects the cell from the extracellular stressors including antifungals. A sturdier cell wall can also serve as a less permeable barrier to antifungal drugs, making them less effective (45). We found that following treatment with caspofungin wild type cells had an increase in the cell wall chitin. This increase was observed at the sub-population level. This was an intriguing finding since sub-population impact of antifungals are a neglected area of study, yet these heterogeneously resistant or tolerant populations may play important roles in antifungal resistance (46–48). Recent studies show that the effect of antifungals at subpopulation level is especially crucial to explain the growth of *Candida* species at supra-MIC concentrations (33). This is mainly due to slow growth of subpopulations compared to the rest of the cells. The heterogeneous nature of axenic microbial cultures, and life in general, is an intricate phenomenon that may significantly contribute to our understanding of antifungal resistance and tolerance. We hope that the single-cell genomics era will enhance our understanding of the antifungal resistance and tolerance pathways in more detail.

Puf4 is a *bona fide* downstream effector of the calcineurin pathway, as it was enriched in a phosphoproteomics screen of the *cna1*Δ mutant (24). Calcineurin was portrayed to be a longstanding player in the antifungal resistance of medically important fungi (49). Many groups have shown that disruption of the calcineurin pathway, genetically or pharmacologically, using novel or repurposed molecules, abolished the calcium homeostasis and led to death. This specific inhibition of calcineurin suggested to us that the calcineurin pathway, especially the downstream effectors, may contain potential targets which can be used as novel antifungal targets (50). Caspofungin treatment causes the translocation of a calcineurin-dependent transcription factor, Crz1. Translocation of Crz1 to the nucleus is an event that induces the transcriptional changes in gene expression in response to caspofungin. Yet, this transcriptional regulation does not contribute to the caspofungin resistance since the *crz1*Δ exhibits a similar caspofungin sensitivity to that of wild type cells (19). On the other hand, the absence of Puf4 causes a hyper-resistant phenotype. While the absolute absence of Puf4 yields a resistance phenotype, we demonstrated that Puf4 protein levels drastically decrease following caspofungin treatment. Interestingly, this suggests that the downregulation of Puf4 may be necessary for the paradoxical resistance, hence the hyper-resistant phenotype in the knockout. The converse was true for the overexpression of Puf4 which showed hyper-sensitivity to caspofungin.

Further work is needed to determine if the phosphorylation state of Puf4 is governed by calcineurin, or if there is another post-translational modification, that is responsible for the rapid reduction in Puf4 protein abundance in response to caspofungin treatment. Inhibition of the mediator of Puf4 protein repression is another potential target to reverse the intrinsic resistance of *C. neoformans* to caspofungin. The post-transcriptional regulation of cell wall homeostasis by Puf4, a calcineurin-regulated RNA-binding protein, is another piece of the regulatory program that results in the intrinsic resistance of *C. neoformans* to caspofungin. Further elucidation of this regulatory program may open new avenues to promote caspofungin sensitivity in *C. neoformans* through adjunctive therapy.

## Materials and Methods

### Yeast strains and molecular cloning

All strains used in this study were derived from *C. neoformans var. grubii* strain H99, a fully-virulent strain gifted from Peter Williamson (UIC, NIAID), which is derived from H99O gifted by John Perfect (Duke University). Primers used to build the knockout and FLAG-tagged complementation strains are included in Table 2. The plasmid construct to establish the PUF4-FLAG-(Hyg) mutant included native promoter and the terminator of the gene. Promoter and the coding sequence were amplified as a single fragment using a forward primer that contained a NotI cut site, and a reverse primer that contained a SalI cut site as well as the FLAG sequence. Puf4 terminator fragment was amplified using forward and reverse primers that contained SalI and BglII sites, respectively. Following restriction digest with respective enzymes, these fragments were cloned into pSL1180 containing the hygromycin B resistance cassette, as described previously (51). Construct containing the PUF4-FLAG was introduced into the *puf4*Δ mutant using biolistics transformation. Copy number were determined using southern blot analysis. A strain that is a single copy tagged complement, and another strain that is a tagged overexpression was established.

### Growth analysis: Spot plates and plate reader assay

Cells were grown overnight at 30°C in 5 mL cultures in YPD broth. Overnight cultures were washed with sterile distilled water and the OD_600_ was equaled to 1 in water. Adjusted cultures were 1:10 serially diluted 5-times and 5 µl of each dilution spotted on YPD agar plates containing indicated concentrations of caspofungin (Sigma). Plates were incubated at 30°C for 3 days and photographed. For the kinetic plate reader assay, overnight cultures were washed with water once and then OD_600_ was equaled to 0.3 in YPD broth. 50 µl of YPD broth containing the 2x the final caspofungin concentration was placed in each well then 50 µl of the OD_600_ adjusted cultures were placed in each well. Plate was incubated at 30°C for 20 hours while shaking in a double orbital fashion and OD_600_ was measured every 10 minutes during this kinetic assay.

### Electrophoretic mobility shift assay (EMSA)

EMSA reactions were set and analyzed as described previously (27). Briefly, all RNA binding reactions contained 5 µg of total protein lysate, 0.5 pmol of the TYE705-labeled oligonucleotide (IDT), and 4 µl 5x EMSA buffer (75 mM HEPES pH 7.4, 200 mM KCl, 25 mM MgCl_2_, 25% glycerol, 5 mM dithiothreitol (DTT), and 0.5 mg/ml yeast tRNA) in a total volume of 20 µl. For competition reactions, 5x, 10x, and 20x more unlabeled wild type or mutant oligonucleotides added in addition to the TYE705-labeled oligonucleotide. Reaction mixtures were incubated at room temperature for 20 minutes, and run on DNA retardation gel, then electrophoresed at 100V. Gels were imaged using a LiCor Odyssey imaging system.

### Motif Search - FIMO: Find Individual Motif Occurrences

Cell wall biosynthesis genes were scanned for the Puf4-binding-element using the FIMO tool on the MEME-suite version 5.1.0 (52). RNA sequences of the cell wall biosynthesis genes in Table 1 were acquired from FungiDB and provided as the input. **UGUA**NNNN**UA** motif was scanned using the default settings. Only the given strand was searched and the p-value criterion was set as p<0.0005 for significance cut-off. Results were manually curated to ensure accuracy in detecting the desired motif.

### RNA stability time-course

Overnight cultures grown at 30°C were used to inoculate 35 mL of YPD broth at the OD_600_ between 0.15 - 0.2 in baffled erlenmeyer flasks. Cultures were grown in baffled flasks at 30 °C while shaking at 250 rpm until they reach the mid-log stage *−* OD_600_ between 0.6 and 0.7. Mid-log stage cultures were supplemented with 250 µg/mL of the transcriptional inhibitor 1,10*−*phenanthroline (Sigma). Then, 5-mL aliquots of each culture were transferred to snap cap tubes and pelleted every 15 minutes for 60 minutes. 50 µl RLT supplemented with 1% *β*-mercaptoethanol was added to each pellet prior to flash freezing in liquid nitrogen. Pellets were stored at −80 °C until RNA extraction. Cells were lysed by bead beating using glass beads. RNA was extracted from each sample using the RNeasy Mini Kit (QIAGEN) following manufacturer’s instructions. RNA was DNase digested on-column using the RNase free DNase Kit (QIAGEN) or using the AMBION TURBO DNA-free Kit (ThermoFisher). cDNA for real time quantitative PCR (RT-qPCR) was synthesized using the Applied Biosystems High Capacity cDNA Reverse Transcription Kit (ThermoFisher) using random hexamers. 800 ng to 1000 ng RNA was used to synthesize cDNA. Samples were quantified using the second-derivative-maximum method and fitted to a standard curve of five 4-fold serial dilutions of cDNA. For experimental samples, cDNA was diluted 1:5 in nuclease-free water. To make the reaction mixture, 5 µl of the 2X SYBR Green Blue Mix (PCRbiosystems) was combined with 4 µl of 1.5 µM primers (970 µl water + 15 µl forward +15 µl reverse). 9 µl reaction mixture was placed in each well and 1 µl of either experimental samples or standards added to their respective wells. Samples from 3 biological samples in duplicate wells were tested. Primer sequences are listed in Table 2 in the along with the gene IDs. Statistical differences were compared by determining the least squares fit of one-phase exponential decay non-linear regression analysis with GraphPad Prism software. Significance between curves was detected with the P-value cut-off of 0.05, which determined that the data from two different curves create different regression lines therefore yielding to different half-lives of the same transcript investigated in different mutants.

### Caspofungin time-course

Cells were grown to the mid-log stage as described in the previous section. At this stage, cultures were supplemented with 16 µg/mL caspofungin and 5 mL aliquots were collected in snap-cap tubes every 15 minutes for 60 minutes. 50 µl of buffer RLT supplemented with 1% *β*-mercaptoethanol was added to each pellet prior to flash freezing in liquid nitrogen. Pellets were stored at −80 °C until RNA extraction. Cells were lysed by bead beating using glass beads. RNA was extracted from each sample, cDNA was synthesized and transcript abundances were calculated using RT-qPCR as described in the previous section. Primer sequences are listed in Table 2. Statistical differences were determined using a two-way-ANOVA.

### Immunoblotting

Cells were grown to the mid-logarithmic stage and half of the culture were treated with 16 µg/mL caspofungin for an hour while the other half was left untreated. Cell pellets were flash frozen in liquid nitrogen and stored at −80 °C. At the time of lysis, 50 µl cold lysis buffer (50 mM Tris HCl pH 7.4, 150 mM NaCl, 1 mM EDTA, 1% Triton X100, 10 µl/ml HALT protease and phosphatase inhibitor [ThermoFisher]) was added and cells were lysed by bead beating using glass beads. 250 µl of the cold lysis buffer was added to the beads and lysate was extracted from the glass beads. Lysate was centrifuged at 20,000 x g for 15 min, and supernatant was transferred to a new tube. Protein quantities were measured using Pierce 660nm Protein Assay Kit (ThermoFisher). 25 µg of protein was run per sample on 4-15% Mini-PROTEAN TGX stain-free precast gels (BioRad) at 150V. Total protein was imaged using a BioRad Gel Documentation System with the stain-free gel setting. Gels were transferred to nitrocellulose membrane and blocked with LiCor Odyssey blocking buffer for an hour. Then, incubated overnight with the mouse Anti-FLAG antibody (1:1000 in Tris-Buffered Saline-Tween20 [TBS-T] with 10% LiCor Odyssey blocking buffer) at 4°C. The blot was washed three times with 15 minutes incubations in TBS-T. Then, LiCor rabbit anti-mouse 800 secondary antibody (1:10000 in TBS-T with 10% LiCor Odyssey blocking buffer) was added. The blot was incubated with the secondary antibody for an hour at room temperature, then washed with TBS-T and blot was imaged using a LiCor Odyssey imaging system.

### Cell wall staining, microscopy and flow cytometry

Cells were grown to the mid-log stage and treated with caspofungin as described previously. Cells were prepared and cell wall components were stained for microscopic and flow cytometric analyses as previously published (43, 53). Briefly, cells were pelleted and washed with 1X PBS once. Cells were fixed with 3.7% formaldehyde for 5 minutes at room temperature, and washed with 1X PBS twice.

Cells were stained with calcofluor and FITC-conjugated wheat germ agglutinin (WGA: Molecular Probes) to visualize chitin. Calcofluor dye stains the total chitin while WGA only stains the exposed chitooligomers. Cells were incubated in the dark with 100 µg/ml FITC-WGA for 35 minutes, then consecutively stained with 25 µg/ml calcofluor white for 15 minutes. Cells were washed twice with 1X PBS before analysis. Stained cells were imaged using a Leica TCS SP8 Confocal Microscope. For microscopy, WGA was detected using the GFP settings and calcofluor was detected using the DAPI settings. Images were taken using the 100X objective. Representative images were shown. Flow cytometry data was acquired using a BD LSRFortessa Cell Analyzer. WGA signal was detected using 488nm laser, and the calcofluor was detected using the 405nm laser. Flow cytometry data were analyzed FlowJo v10.0 software. Representative histogram graphs were shown.

Cells were stained with Aniline Blue to detect the *β*-1,3-glucan levels. Unfixed mid-log stage cells in YPD, untreated and treated with 16 µg/ml caspofungin, were washed with 1X PBS and stained with 0.05% Aniline Blue (Wako Chemicals, Japan) for 10 minutes, and then cells were imaged using the DAPI channel on the Leica TCS SP8 Confocal Microscope. Microscopy images were analyzed using the Fiji (Image J, NIH). Mean fluorescent intensity of at least 100 cells in 3-4 different fields were quantified and normalized. Representative images are shown.

### Statistical Analysis

Data analysis was performed using the GraphPad Prism software version 6. Each figure legend contains the statistical test information that is used to assess the statistical significance. Briefly, we have utilized the one-phase exponential decay analysis to determine the half-life of the mRNAs analyzed. Immunoblot data was analyzed using unpaired t-test with Welch’s correction. Gene expression and microscopy quantification data were analyzed using either one-way or two-way-ANOVA followed by a post-hoc multiple comparison test. For all of the graphs, *: p<0.05, *** p<0.001 and ****: p<0.0001. All error bars represent the SEM.

## Acknowledgements

We would like to acknowledge Dr. Amanda L. M. Bloom for advice and stimulating discussions. This work was funded by NIH R21 AI133133 to JCP.

